# RNA modifications detection by comparative Nanopore direct RNA sequencing

**DOI:** 10.1101/843136

**Authors:** Adrien Leger, Paulo P. Amaral, Luca Pandolfini, Charlotte Capitanchik, Federica Capraro, Isaia Barbieri, Valentina Migliori, Nicholas M. Luscombe, Anton J Enright, Konstantinos Tzelepis, Jernej Ule, Tomas Fitzgerald, Ewan Birney, Tommaso Leonardi, Tony Kouzarides

**Author notes:** Co-first authors (Adrien Leger, Paulo P. Amaral). Co-last authors. Correspondence.

## Abstract

RNA molecules undergo a vast array of chemical post-transcriptional modifications (PTMs) that can affect their structure and interaction properties. To date, over 150 naturally occurring PTMs have been identified, however the overwhelming majority of their functions remain elusive. In recent years, a small number of PTMs have been successfully mapped to the transcriptome using experimental approaches relying on high-throughput sequencing. Oxford Nanopore direct-RNA sequencing (DRS) technology has been shown to be sensitive to RNA modifications. We developed and validated Nanocompore, a robust analytical framework to evaluate the presence of modifications in DRS data. To do so, we compare an RNA sample of interest against a non-modified control sample. Our strategy does not require a training set and allows the use of replicates to model biological variability. Here, we demonstrate the ability of Nanocompore to detect RNA modifications at single-molecule resolution in human polyA+ RNAs, as well as in targeted non-coding RNAs. Our results correlate well with orthogonal methods, confirm previous observations on the distribution of N6-methyladenosine sites and provide novel insights into the distribution of RNA modifications in the coding and non-coding transcriptomes. The latest version of Nanocompore can be obtained at https://github.com/tleonardi/nanocompore.

## 1 Introduction

RNA post-transcriptional modifications (PTMs) are a pervasive feature common to all domains of life. They arise from covalent alteration or isomerisation of nucleotides, typically involving the addition of chemical groups to different positions of the nitrogenous bases or the ribose cycle. To date, over 150 modifications have been found throughout all classes of RNAs, with the most common modification being methylation^1^. PTMs are deposited and catalytically removed by specific enzymes and can be recognized by specific ‘reader’ proteins. Overall, PTMs influence fundamental properties and functions of RNAs, including their stability, structure, intermolecular interactions and cellular localization^2,3^. Over the past few decades, methods such as RNA fragmentation and liquid chromatography combined with mass spectrometry (LC-MS) have provided an extensive characterisation of PTMs in the abundant classes of RNA, especially tRNAs, rRNAs and snoRNAs^4^. More recently, various strategies harnessing high-throughput sequencing have allowed detection and mapping of PTMs in the less abundant messenger RNAs (mRNAs) and regulatory noncoding RNAs (ncRNAs)^5,6^. At the same time, enzymes catalysing the deposition of PTMs in mRNAs have been identified at a steady pace, allowing mechanistic studies into the role of PTMs in post-transcriptional regulation as well as their impact on cancer and developmental diseases^7–11^. N6-Methyladenosine (m6A) is the best characterised PTM and the most abundant in mRNAs and long noncoding RNAs (lncRNAs). It is deposited mainly by the METTL3/METTL14/WTAP complex and has a variety of functions such as regulation of nuclear export, translation and degradation of RNAs. m6A is found preferentially at the consensus motif DRACH (D=A, G or U; R=A or G; H=A, C or U) and is highly enriched near mRNA stop codons^12,13^. Pseudouridine is another well characterised modification, occurring via an isomerisation of uracil catalysed by enzymes of the pseudouridine synthase (PUS) family and the dyskerin pseudouridine synthase 1 (DKC1) enzyme^14^. It is the most abundant internal PTM present in RNAs, including the highly conserved transcriptional regulator 7SK RNA^15^. The majority of current methods for mapping PTMs rely on RNA immunoprecipitation, chemoselective alteration or specific signatures resulting from reverse transcription (RT), and therefore suffer from the shortcomings of these approaches, including (1) the need to develop *ad hoc* protocols for each PTM, (2) cross reactivity or low sensitivity of antibodies or chemical reactions and (3) biases induced by the complex multi-step experimental protocols^5,6^.

The recent advances in Nanopore direct RNA sequencing (DRS) have allowed, for the first time, direct sequencing of full-length native RNA molecules without the need for RT or amplification. Importantly, a number of studies have shown that DRS data intrinsically contain information about RNA modifications^16–18^, and the development of reliable tools to extract this information remains a significant and immediate challenge. In Nanopore DRS a single DNA or RNA molecule is ratcheted by a molecular motor through a protein pore embedded in a synthetic membrane. The passage of nucleobases through the narrowest section of the pore (reader-head) alters the flow of ions across the membrane, depending on the chemical composition of the bases. At any given point in time approximately 5 nucleotides reside within the reader-head of R9 pores, leading to a strong 5-mer specific signal alteration. Crucially, the presence of nucleotide modifications can induce discernible shifts in current intensity and in the time the nucleic acid sequence resides inside the pore (dwell time)^16^. These parameters can be used to train predictive models capable of identifying modified sites at single nucleotide resolution from raw current signal. This type of strategy has been successfully applied by several academic groups^19,20^ as well as by Oxford Nanopore Technologies (ONT) (e.g. tools such as Megalodon and Guppy) to detect DNA methylation. However, creating training sets and predictive models for RNA modifications has so far proven to be a much more challenging task. There are three notable pieces of software currently available for RNA modifications detection: Tombo^21^, EpiNano^22^ and MINES^23^. Tombo is a toolset developed by ONT for detection of DNA modifications, which was later extended to RNA but only with moderate success. The TOMBO RNA analysis framework has several unresolved issues, surrounding RNA signal alignment and ONT recently announced their intention to move towards different strategies (London Calling 2019). On the other hand, EpiNano mostly relies on a model of basecalling errors induced by the presence of m6A. This model was generated from a training set obtained by *in vitro* transcription of RNAs in the presence of either canonical A or m6A. Although powerful, this approach is potentially affected by the limited number of sequence contexts that contain m6A (i.e. limited kmer coverage) and is difficult to generalise to other modifications. MINES is a classifier trained on m6A canonical motif sites identified by miCLIP. This approach is innovative and offers isoform resolution, but is limited to m6A sites within 4 specific DRACH sequences and could be affected by the same biases and/or limitations as miCLIP. There is therefore a currently unmet need for a robust method capable of reliably detecting multiple RNA modifications in any possible sequence contexts.

Here we introduce Nanocompore, a flexible and versatile analysis method dedicated to the detection of RNA modification from DRS datasets. To identify potential modification sites, Nanocompore uses a model-free comparative approach where an experimental RNA sample is compared against a sample with fewer or no modifications. Potentially, this can be applied to any modification, provided that an appropriate control sample strongly depleted of the modification is available. Nanocompore includes several unique features: (1) robust signal realignment based on Nanopolish, (2) modelling of the biological variability, (3) ability to run multiple statistical tests, (4) prediction of RNA modifications using both signal intensity and duration (dwell time) and (5) availability of an automated pipeline that runs all preprocessing steps. Finally, the results generated by Nanocompore can also be leveraged to infer RNA modifications at single molecule resolution.

We validated the performance and sensitivity of Nanocompore using multiple strategies. Specifically, we tested our method on hundreds of *in silico* generated datasets as well as a dataset generated from an *in vitro* modified oligonucleotide sample. We then applied Nanocompore to the detection of METTL3-dependent m6A sites in the human polyA+ transcriptome as well as in targeted ncRNAs. Our findings were cross-validated with methylated individual-nucleotide-resolution UV cross-linking and immunoprecipitation (miCLIP). Finally, using an *in vitro* transcribed (IVT) control sample, we showed the ability of Nanocompore to map large numbers of candidate sites, albeit without identifying which modification.

## 2 Results

### 2.1 Nanocompore data preparation and statistical basis

Nanocompore detects potential RNA modifications by comparing DRS datasets from one experimental test condition containing specific RNA modifications to one control condition containing significantly fewer or no modifications. Ideally, the control RNA is isolated from a cell harbouring either a knock-down (KD) or a knock-out (KO) of a gene encoding an RNA modifying enzyme. Alternatively, for small scale comparison, it is also possible to use either an *in vitro* transcribed or synthetic RNA containing canonical RNA bases only. Before running Nanocompore, the DRS data need to be preprocessed. These steps can be effectively and easily streamlined using our companion Nextflow pipeline (https://github.com/tleonardi/nanocompore_pipeline) that automatically runs the entire analysis starting from raw Nanopore data (detailed in **Figure 1A**). Brefly, raw fast5 reads from all input DRS datasets are basecalled and quality control is performed to assess the consistency of the datasets^24^. Basecalled reads are then aligned to a reference transcriptome. Alignment information is used by the robust Nanopolish algorithm^19^ to realign raw nanopore signal to the expected reference sequence. This step is essential for annotating and slicing the signal corresponding to each kmer sequenced. Finally, these data are collapsed by kmer and indexed by read. The resulting realigned reads are then processed with Nanocompore (https://github.com/tleonardi/nanocompore) to identify modified bases (**Figure 1B**). Firstly, reads are grouped by reference transcript and transcripts with coverage above a user-specified threshold are used for subsequent analyses. Then, two parameters - the median signal intensity and the log_10_ (dwell time) - are collected from each read and aggregated at the transcript position level. These descriptive data are stored on disk to allow users to visually inspect the modification-induced effect on the signal. The aggregated data are compared in a pairwise fashion, one position at the time. For the identification of modified positions, Nanocompore currently supports 3 robust univariate pairwise tests: Kolmogorov-Smirnov (KS), Mann-Whitney (MW) and Welch’s t-test. In addition, we also implemented a more advanced bivariate classification method based on 2 component Gaussian mixture model (GMM) clustering followed by a statistical test to determine if there is a significant difference in the distribution of reads into the two clusters between conditions. This test is implemented either as a logistic regression or an ANOVA test. Furthermore, we and others observed that DNA and RNA modifications can have an intrinsic effect on the local signal upstream or downstream of the modification position. Thus, to evaluate the effect of modifications on the proximal sequence context, we implemented an established statistical method that combines the non-independent p-values to produce a combined p-value representative of multiple neighbouring kmers^25^. The p-values are then corrected for multiple tests using Benjamini-Hochberg’s procedure^26^ and the results are stored in a lightweight database. Users can obtain a tabular text dump of the database or use the extensive Nanocompore API to explore the results and generate ready-to-publish plots.

**Figure 1:**
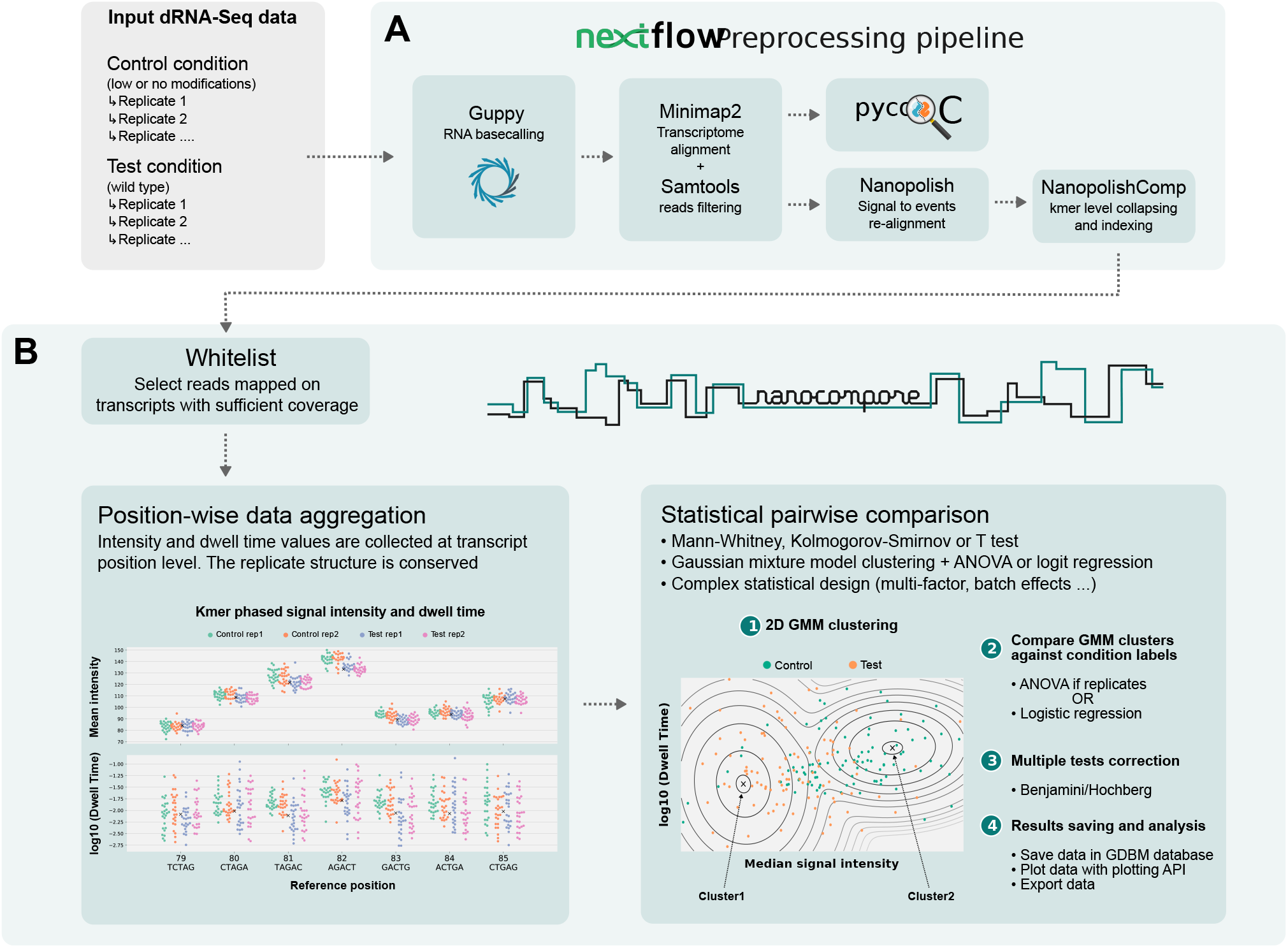
Overview of data preparation and Nanocompore steps. **A:** Raw fast5 reads from 2 conditions are basecalled with Guppy, filtered with Samtools and the signal is then resquiggled with Nanopolish eventalign. The output of Nanopolish is then collapsed and indexed at the kmer level by NanopolishComp Eventalign_collapse. **B:** Nanocompore aggregates median intensity and dwell time at transcript position level. The data is compared in a pairwise fashion position per position using univariate tests (KS, MW, t-tests) and/or a bivariate GMM classification method. The p-values are corrected for multiple tests and these data are saved in a database for further analyses.

### 2.2 *In silico* and *in vitro* validation

In order to estimate the sensitivity and specificity of Nanocompore, we first generated an unmodified RNA model from an *in vitro* transcribed (IVT) RNA DRS dataset containing A, U, C and G canonical bases only^17^. For each 5-mer, we estimated the distribution type and parameters that best fit the observed current intensity and dwell time values. Based on this model, we simulated a reference *in silico* dataset from a semirandomly generated unmodified reference sequence (see Materials and Methods). To mimic signal changes induced by RNA modifications we then *in silico* generated 144 “modified” datasets where the dwell time and signal intensity at defined positions are shifted from the means of the unmodified model by a varying multiple of standard deviations. For each combination of dwell time and signal intensity shift we generated 6 datasets with varying fractions of modified reads (ranging from 10% to 100%) (**Figure 2A**). We ran Nanocompore on each dataset to test its sensitivity and specificity for identifying these known modified positions. We observed that all the statistical tests implemented in Nanocompore had near perfect precision and recall (0.9889 ≤ AUROC ≤ 0.9947) when the simulated shifts from the model were greater than 2 standard deviations (SD) for both intensity and dwell time and the fraction of modified reads was above 25% (**Figure 2B** and **Sup. Fig. S1**, page 22). The GMM method performed better than other methods when only as little as 25% of the reads were modified, although requiring a relatively large shift between the 2 populations (≥ 2 SD). On the other hand, the KS tests achieved near perfect precision and recall with milder intensity or dwell time shifts (1 SD) at the expense of the need for a larger population of modified reads. (**Sup. Fig. S1**, page 22). For these reasons, Nanocompore implements both tests and users can choose which one to use depending on their experimental settings.

**Figure 2:**
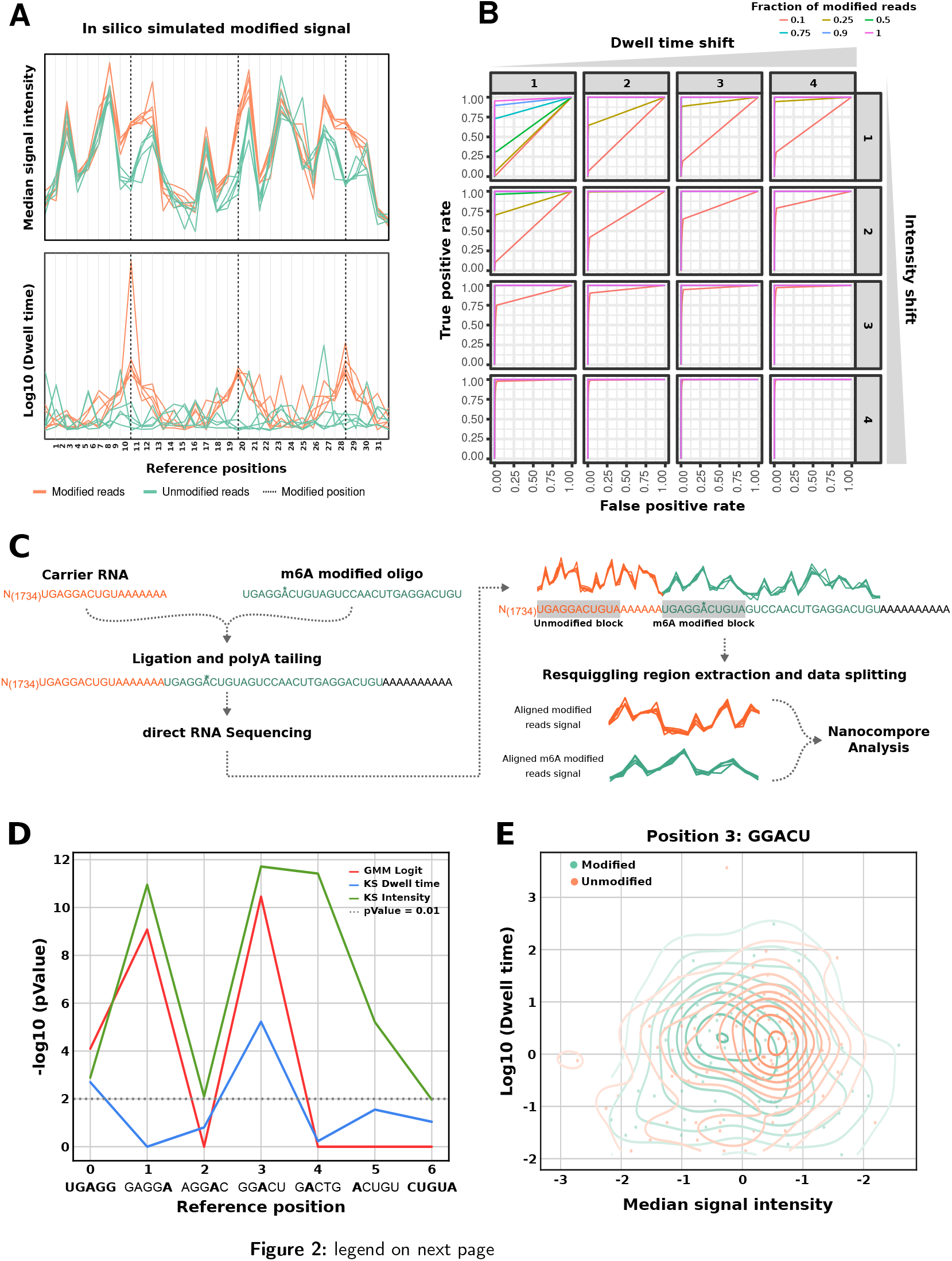
*In silico* and *in vitro* validation of Nanocompore. **A:** Position plots showing the median intensity (top) and dwell time (bottom) for simulated data generated by Nanocompore SimReads. Dashed vertical lines indicate modified positions, where a clear signal shift between unmodified (green) and modified (orange) reads can be seen. **B:** ROC curves obtained with the GMM method on *in silico* generated data. The different subplots indicate varying amounts of intensity and dwell time shifts from the unmodified model (from 1 to 4 standard deviations), whereas the different colors indicate varying fractions of modified reads in the generated data (from 10% to 100%). All the comparisons are against a fully unmodified reference dataset. **C:** Diagram depicting the experimental and signal processing workflow used to run Nanocompore on a synthetic RNA carrying an N6-methyladenosine residue. Briefly, the RNA sequence was obtained by ligating an oligo carrying an m6A residue at position 6 to a longer unmodified carrier obtained by IVT of the firefly luciferase gene. The data obtained by DRS was parsed to extract the signal from the 2 short regions corresponding to a “UGAGGACUGUA” with either an A or an m6A in the middle. **D:** Plot showing the Nanocompore p-values (-log_10_, y-axis. Tests: GMM+logit and KS) for the synthetically modified GGACU kmer and those surrounding it. **E:** Scatter plot showing the median signal intensity (x-axis) and scaled dwell time (log_10_, y-axis) for the synthetically modified kmer. The concentric lines show the kernel density estimates for the two samples.

As a further control for Nanocompore sensitivity, we re-analysed a DRS dataset generated by the Nanopore RNA Consortium consisting of a synthetic oligonucleotide carrying m6A at a known position^17^ (**Figure 2C**). The data were preprocessed as described before and then split into a modified and an unmodified subset (**Figure 2C** and Materials and Methods). We found that Nanocompore correctly identified the modified position as highly significant (p-values of 3.50×10^-11^, 5.95×10^-6^ and 1.93×10^-12^ for the GMM+logit, KS_dwell and KS_intensity test respectively, **Figure 2D**), with the clear formation of 2 clusters for reads corresponding to the modified and unmodified data (**Figure 2E**). We also observed that the intensity shift spreads to the adjacent kmers containing the m6A residue (positions 1 to 5, **Figure 2D**). This shows that a modification can alter the signal locally and supports the rationale of combining the p-values of neighbouring kmers.

### 2.3 Transcriptome-wide m6A profiling

Having validated the accuracy of Nanocompore on simulated and synthetic data, we sought to apply it to map m6A in a biologically relevant context. METTL3-METTL14 heterodimers form a N6-methyltransferase complex that methylates adenosine residues at the N(6) position of specific RNAs. Since m6A is required for development and maintenance of acute myeloid leukemia^8,27^, it is of particular importance to accurately map it in leukemia cells. We therefore used DRS to profile the poly-A+ transcriptome of human MOLM13 cells with inducible shRNA-mediated knockdown (KD) of METTL3, as well as control Wild Type (WT) MOLM13 transfected with a scrambled shRNA. We sequenced RNA from two biological replicates per condition on independent Minion flow cells after 4 days of induced KD of METTL3, yielding a total of 3,768,380 reads. After applying a stringent 30X coverage threshold, we obtained data for 752 unique transcripts robustly expressed in all samples (**Sup. Fig. S2A-C**, page 24). Overall, we observed a high correlation of expression levels between samples showing the consistency of the datasets (R2 of 0.969, **Sup. Fig. S2D-F**, page 24). We then used Nanocompore to map the location of METTL3-dependent m6A sites in human transcripts from MOLM13 cells. Applying a false discovery rate of 1% (GMM+logit method, see Materials and Methods) we found 6,021 significant kmers in 437 distinct transcripts, which could be grouped into 4,094 clusters (clustering distance 5nt) averaging 9.37 clusters per transcript (**Figure 3A,B**). As an example, we found 61 significant clusters (83 kmers with p-value<0.01, **Figure 3C**) in the β-actin (ACTB, ENST00000646664) mRNA. Interestingly, the 3 most significant β-actin hits are “GGACU” kmers, perfectly matching the canonical m6A DRACH motif (**Figure 3D-F**). On a transcriptome-wide scale, we reproduced previous observations showing that METTL3-dependent m6A sites are enriched in the immediate vicinity of mRNA stop-codons (**Figure 3G**)^12,28^. Additionally, we used Sylamer^29^ to identify enriched kmers in the Nanocompore significant sites, finding a 4.3 fold enrichment for the consensus GGACU motif in the Nanocompore sites with p-value<0.01 (hypergeometric p-value=4.3×10^-21^ **Figure 3H**). Lastly, we generated miCLIP datasets from MOLM13 cells targeted with METTL3 CRISPR gRNAs to compare the results obtained with Nanocompore with an orthogonal high-resolution method. Nanocompore positive sites show a peak of normalized miCLIP crosslink sites in WT cells, whereas it is significantly reduced in METTL3 KO cells (log2 fold change −0.45, p-value=7.73×10^-62^, Mann-Whitney test, **Figure 3I** and **Sup. Fig. S3**, page 26). Overall, these results show that Nanocompore is capable of identifying enzyme-specific RNA modifications transcriptome-wide and that these findings are in agreement with previous techniques.

**Figure 3:**
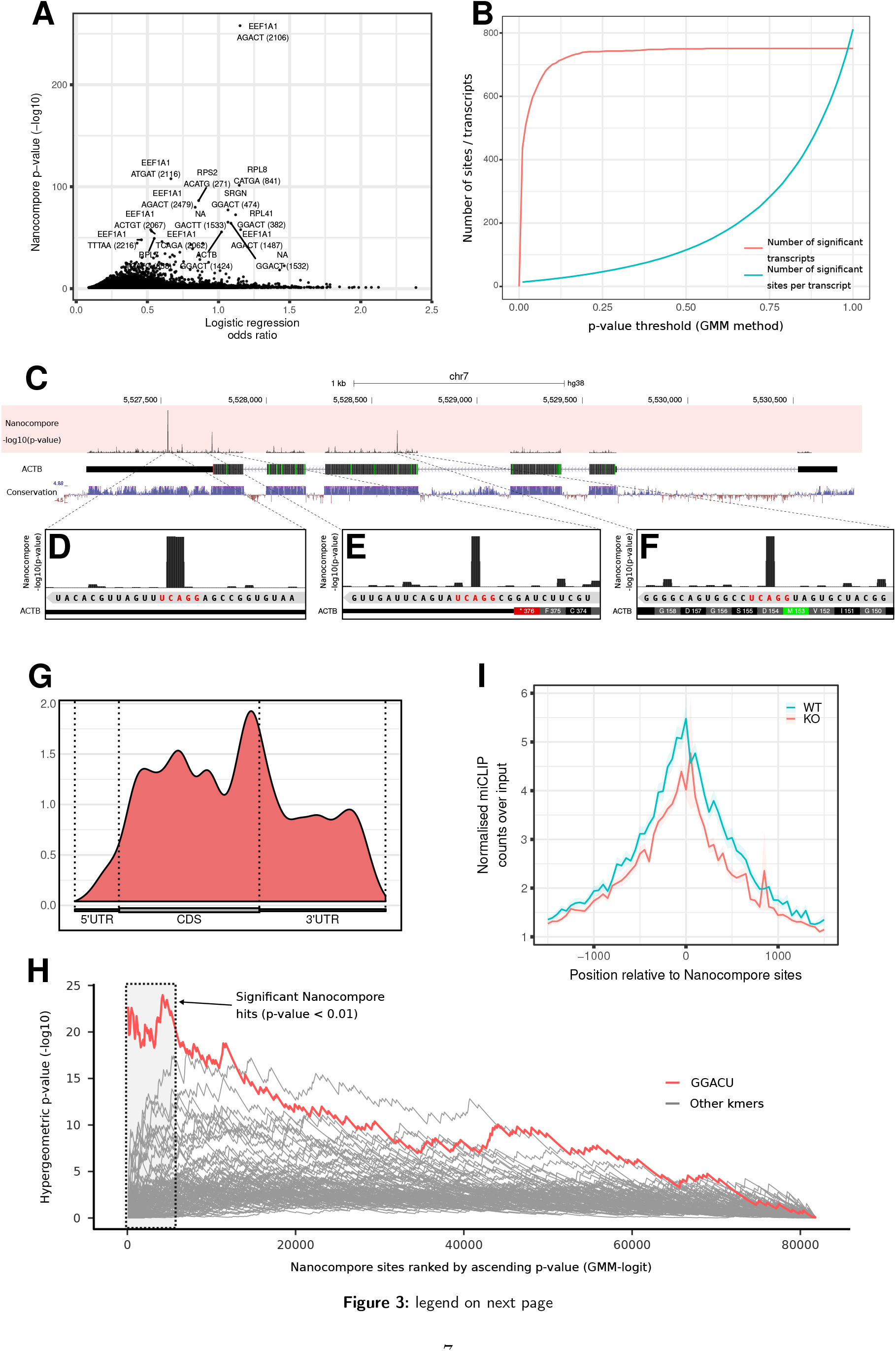
m6A profiling in MOLM13 cells. **A:** Sharkfin plot showing the absolute value of the Nanocompore logistic regression log odd ratio (x-axis) plotted against its p-value (-log10, y-axis, see Material and Methods). Each point represents a specific kmer of a transcript. **B:** Red line, total number of transcripts with at least one significant Nanocompore site at varying p-value thresholds (x-axis). Blue line: average number of significant Nanocompore sites per transcripts at varying p-value thresholds. **C:** Genome browser screenshot showing METTL3-dependent m6A sites in the ACTB transcript. The p-value track reports the Nanocompore GMM+Logistic regression method (see Material and Methods). **D-F:** As in C but showing the three most significant β-actin sites at higher magnification. The sequence reported at the bottom corresponds to the RNA sequence in the 3’ to 5’ orientation, as the ACTB transcript is encoded on the minus strand. The overlapped m6A consensus GGACU sequences are highlighted in red. **G:** Metagene plot showing the coverage of m6A sites across transcripts. **H:** Sylamer plot showing kmer enrichment in Nanocompore significant sites. The x-axis reports all Nanocompore sites with p-value<0.5 ranked from the most to the least significant. The y-axis reports the Sylamer hypergeometric p-value of enrichment of a certain motif in the first x Nanocompore sites vs the rest. The vertical dotted line delineates Nanocompore sites with p-value<0.01 (to the left of the line). **I:** m6A miCLIP coverage of clusters of significant Nanocompore sites (GMM logit p-value<0.01). Coverage calculated in transcriptome-space. The y-axis shows the mean input-normalised signal across sites of the average coverage (counts per million) in four and two replicates for WT and KO respectively. The shaded area shows the standard error of the mean across sites. The difference between WT and KO in the windows 0+/-100nt is statistically significant (p-value=7.73×10^-62^, Mann-Whitney test).

The identification of RNA modifications outlined so far operates at consensus level, i.e. looking at the distribution of signal across the entire population of reads. However, the information obtained from GMM clustering at the population level can be leveraged to calculate the probability of each read to belong to the modified or unmodified cluster. Hence, it is possible to assign modification probabilities at the single-molecule, single-site level. As a proof of concept, we calculated the single-molecule modification probabilities of the three β-actin high-confidence m6A sites previously described (**Figure 3C-F**). We found that these three sites are methylated at different degrees: 45% of β-actin molecules methylated with high-confidence (probability >0.75) at A652, 23% at A1324 and 49% at A1535. As expected, we also found that the fraction of methylated molecules decreased at all three sites in the METTL3 KD condition (26%, 14% and 27% of molecules methylated at A652, A1324 and A1535 respectively, **Figure 4A-C**). We further asked whether the presence of an m6A modification at one of these three sites influences the probability that the same molecule is modified at the other sites. Taking into account the underlying frequency of modification at each site, we calculated the conditional probabilities for all possible combinations of 0, 1, 2 or 3 modifications to co-occur in the same molecule (**Figure 4D**). This analysis showed that the observed and expected modification frequencies do not differ significantly, suggesting that methylation of these three sites are independent events (p-value=0.4, see Materials and Methods).

**Figure 4:**
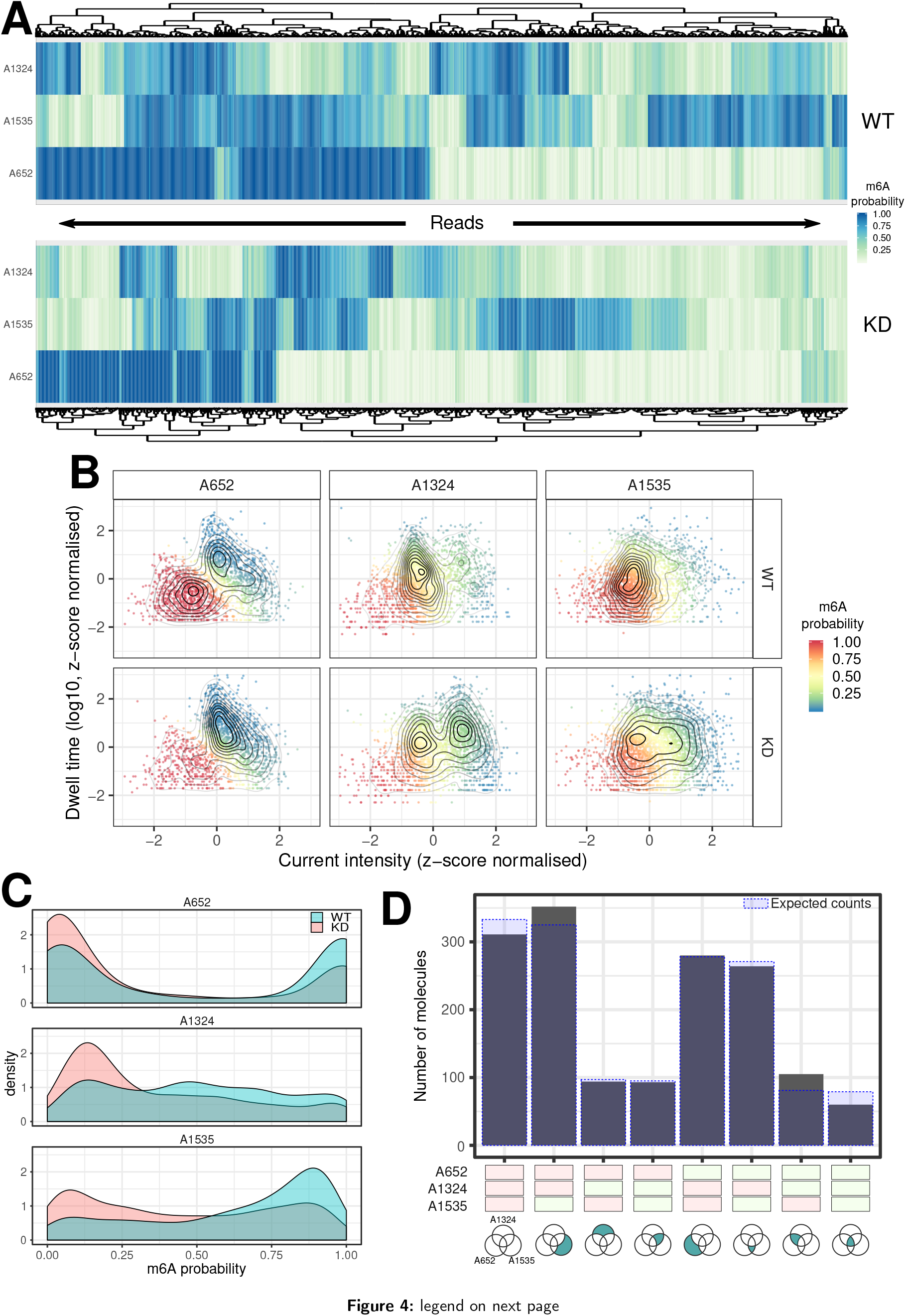
Single molecule identification of m6A sites. **A:** Heatmap organised by hierarchical clustering showing the probability of A652, A1324 and A1535 in the β-actin gene to be modified in the WT and METTL3 KD samples. Each column corresponds to a single molecule. **B:** Scatter plot with overlaid kernel density estimates showing the scaled median intensity vs the scaled log_10_ dwell time for each read covering A652, A1324 and A1535. Data points are color coded according to the probability that the read belongs to the cluster of m6A modified reads. For visualisation purposes the x- and y-axis were limited to the +/-3 range. **C:** Density plot showing the distribution of modification probability for A652, A1324 and A1535 of β-actin in WT (blue) and KD (red). **D:** Bar chart showing the number of molecules identified in each of the 8 possible m6A configurations for the A652, A1324 and A1535 sites of β-actin. Each site was considered modified if the modification probability was >0.75. The shaded blue areas indicate the expected number of molecules in each given configuration under the null hypothesis of independence of the three modifications.

### 2.4 Modification mapping in snRNA 7SK by high coverage targeted sequencing

Using the same inducible METTL3 KD and control cells as above, we next performed high-coverage targeted DRS of the human non-coding snRNA 7SK. To do so, we designed a custom nanopore sequencing adapter targeting the 3’ end of 7SK (see Materials and Methods and **Table S2**). With this approach we achieved consistently high coverage in all the samples (average of 4,844 reads per sample). 7SK is a highly structured RNA with numerous binding sites for interacting proteins, which together form the 7SK snRNPs (**Figure 5A**). Nanocompore analysis of 7SK in METTL3 KD cells identified 24 significant kmers across its entire sequence (p-value<0.01, **Figure 5A,B**). The most significant hit falls in the UGAUC kmer at position 41 (**Figure 5A-D**), which corresponds to the 5’ palindrome of the double-stranded and structurally conserved binding site for HEXIM1^30^. Interestingly, the 3’ GAUC palindrome at position 64 is also a significant site (**Figure 5A,B,D**). These results suggest that the two central adenosines of the double stranded HEXIM1 binding site (A43 and A65) are both methylated by METTL3. We also identified 5 significant overlapping kmers between positions 229 and 250 in the terminal loop of hairpin 3 (HP3) (**Figure 5A,B**). This region was recently shown to be the binding site for RNA-binding motif protein 7 (RBM7), which mediates the activation of P-TEFb by releasing it from 7SK snRNP^31^.

**Figure 5:**
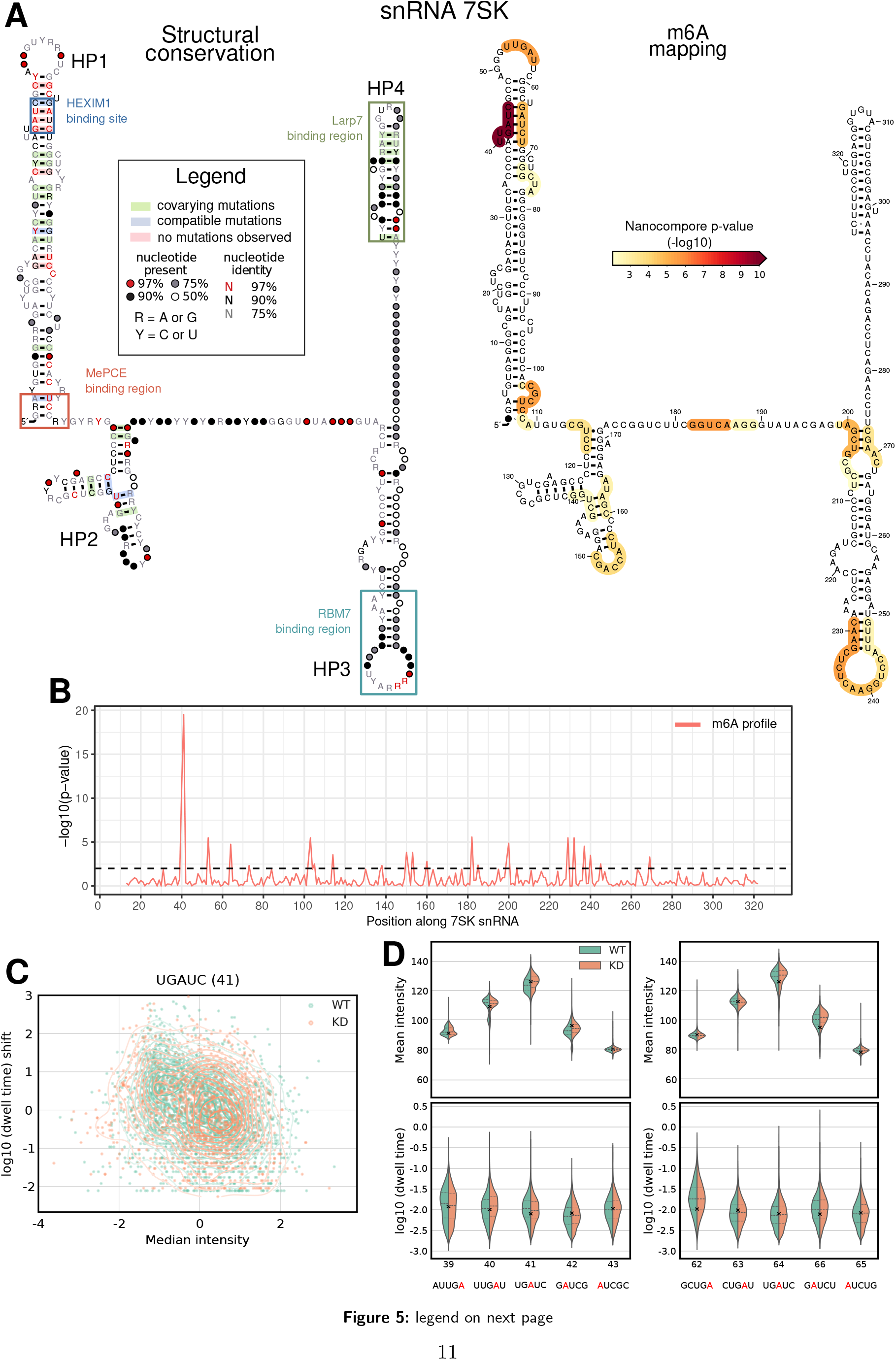
m6A identification in 7SK RNA. **A:** On the left, secondary structure of 7SK showing positions of known protein binding sites and structural conservation. On the right, secondary structure of 7SK with the Nanocompore p-value (METTL3-KD vs WT, GMM-logit) overlaid as a color scale. For each nucleotide the color indicates the lowest p-value among those of the 5 kmers that overlap it. Only p-values<0.01 are shown in color. **B:** m6A profile of 7SK, showing the Nanocompore GMM-logit p-value (y axis, −log10) across the transcript length. **C:** Scatter plot showing the scaled median intensity vs the scaled log10 dwell time for each read covering kmer 41 of 7SK. Each point shows data for a distinct read color coded according to the sample. The contour lines show the kernel density estimates for the two samples. For visualisation purposes the x- and y-axis are truncated at −4 and +3 respectively. **D:** Violin plots showing the distributions of median intensity (top) and scaled log10 dwell time (bottom) for the Hexim1 binding sites and neighbouring kmers. All coordinates refer to the first nucleotide of each kmer relative to ENST00000636484.

We next sought to extend our investigation of 7SK to include other modifications in addition to m6A. To this end we used IVT to generate large amounts of 7SK RNA devoid of all modifications. We then sequenced this IVT 7SK by DRS and analysed the resulting data with Nanocompore, using the dataset from targeted sequencing of WT MOLM13 cells as the reference condition. This approach potentially allows mapping of all RNA modifications in targeted RNAs, albeit without revealing the type of each modification. We identified 68 significant kmers spread across the entire 7SK sequence (1% FDR, **Sup. Fig. S4A**, page 27). The most significant region identified is ~10nt long and is located at the stem-loop boundary of HP3 (**Sup. Fig. S4B**, page 27). This region encompasses the m6A site identified at position A245 by the analysis of METTL3-KD, as well as a known pseudouridine site at position U250 (**Sup. Fig. S4C**, page 27)^15^. We also observed a significant change between IVT and WT RNA samples at A43 (UGAUC kmer, p-value=0.0608) and A65 (GCUGA and CUGAU kmers, p-values=0.0839 and 0.0346, respectively), supporting the presence of the two m6A sites that we identified above in the double stranded HEXIM1 binding site. To confirm that the signal observed at U250 of 7SK is indeed due to the presence of pseudouridine at this position, we repeated targeted DRS after knocking down dyskerin pseudouridine synthase 1 (DKC1), the enzyme responsible for catalysing pseudouridine formation at this position^15^. Nanocompore analysis of these data revealed a significant kmer at position 317 (AGUCU, **Sup. Fig. S5A**, page 28), indicating U318 or U320 as potential DKC1-mediated pseudouridine sites. However, U250 did not pass the FDR threshold, but we did find three significant kmers covering the adjacent stretch of three U at positions 246-248 (**Sup. Fig. S5A**, page 28, kmers UCCAU, CCAUU and AUUUG, KS test on current intensity). In this experiment we also simultaneously targeted 3 other ncRNAs; RMRP, RPPH1 and U2 snRNA. We achieved sufficient coverage for all of them and found 6 significant kmers for RMRP and 2 for U2 (GMM p-value<0.01, **Sup. Fig. S5B-D**, page 28).

## 3 Discussion and conclusions

In recent years substantial progress has been made in our understanding of the roles and functions of RNA PTMs. In fact, it is becoming increasingly clear that they can affect RNA function and metabolism. The diverse range of RNA PTMs biological roles are mediated by their capacity to dynamically regulate the physical and chemical properties of RNA molecules, for example by creating or masking binding sites, altering RNA structure or modulating expression and subcellular distributions^7,11^. However, fully understanding the breadth and scope of RNA modifications as well as their dynamic regulation in physiological and pathological context requires efficient and accurate methods to detect their presence and to map them to the respective RNA sequence contexts.

In this paper we introduce Nanocompore, a robust method for the identification of any RNA modifications site from Nanopore DRS data. Nanocompore performs a signal level comparison between two conditions, allowing identification of significant changes indicative of the presence/absence of RNA modifications (**Figure 1**). Our approach has several advantages over alternative RNA PTM mapping methods. First, it is based on Nanopore DRS which means it is not affected by retro-transcription or PCR amplification mediated biases that other genome-wide strategies are subject to. Second, it maps RNA modifications in the context of long reads, giving critical information on the impact of RNA PTMs on individual gene isoforms. Third, our comparative strategy does not require any training and can be applied as-is to any RNA modification, as long as a modification-depleted reference sample is available. Fourth, Nanocompore is the first generalist method which can be used to map physically linked RNA modifications at single molecule resolution. Finally, we implemented a pipeline in the Nextflow Domain Specific Language that automatically runs all processing steps, from the raw data up to the execution of Nanocompore, thus greatly simplifying the bioinformatics work. As previously mentioned, there are several alternatives offering some the functionalities included in Nanocompore, such as Tombo^21^, EpiNano^22^ and MINES^23^. However, they target specific modifications or use simpler statistical frameworks making them hard to compare. Given the unique strengths and pitfalls of each method, we would encourage users to test other tools and integrate the results with Nanocompore predictions.

We validated the performance of Nanocompore using a purposely generated *in silico* dataset as well as an *in vitro* control oligonucleotide (**Figure 2** and **Sup. Fig. S1**, page 22). We then applied Nanocompore to a biologically relevant dataset, where the main m6A writer enzyme (METTL3) was knocked-down in human cells (**Figure 3**, **Sup. Fig. S2**, page 24 and **Sup. Fig. S3**, page 26). In this context, we were able to recapitulate previous observations and provide novel insights. For example, we found m6A to be enriched toward mRNA stop codons as well as for the short motif DRACH. Furthermore, we could also confirm by miCLIP that m6A is enriched at the sites identified by Nanocompore. This experiment shows the feasibility of using Nanocompore to detect RNA modifications transcriptome-wide, even at relatively shallow sequencing depths. As an additional proof-of-concept, we performed high coverage targeted sequencing of non-polyadenylated ncRNAs, identifying multiple putative modification sites in the 7SK snRNA (**Figure 5** and **Sup. Fig. S4**, page 27). In addition to METTL3-dependant m6A sites we were also able to profile Dyskerin-dependent Pseudouridine candidate sites as well as the overall modification landscape by comparing our sample with an IVT 7SK control (**Sup. Fig. S5**, page 28).

We showed that Nanocompore performs well with mixed populations of modified/unmodified reads in the control and experimental samples. Although it is currently unsuitable for the identification of very low-frequency modifications, our *in silico* benchmarks show good sensitivity in mixed populations where only 25% of reads are modified. This observation strengthens the importance of having good control conditions, such as high efficiency knock-downs, knock-outs or IVT samples.

An additional feature of Nanocompore is that by analysing knock-down or knock-out samples it intrinsically assigns RNA modifications to their specific writer enzymes, thus allowing one to discern the individual roles of multiple enzymes that catalyse the same modification. However, an important caveat to be considered when pursuing this approach - as well as any other method based on loss-of-function of catalytic enzymes - is that it is likely that some RNA modifications are catalysed by multiple enzymes with partially overlapping functions that could compensate each other when individually depleted. A further complication is the potential interaction of neighbouring modifications: for example, one modification might be required in order for another (possibly different) one to be deposited or removed. Then, the loss of one enzyme could indirectly result in signal differences on bases harbouring a different modification, which depends on the one catalysed by the enzyme. These possible scenarios and secondary effects may be confounding factors for Nanocompore analysis and currently cannot be accurately resolved with our method.

In the field of epigenetics, CpG methylated sites have long been known to influence each other and methylation can quickly spread to adjacent CpGs^32^. On the other hand the interplay between RNA modifications is still widely unknown. Hence, in addition to isoform-level resolution of RNA modifications, it is becoming increasingly important to obtain information about modification stoichiometry and combinatorics. One of the major advantages of Nanocompore is its ability to detect RNA modifications at single molecule resolution. We applied our analysis to the most significant m6A sites found by Nanocompore in β-actin mRNA and found that multiple methylated residues are present in the same molecule independently of one another at a given time. These observations suggest the presence of highly site-selective intramolecular deposition and/or removal of m6A. This is the first observation of this kind to date, and it will need to be cross-validated when other methods enabling the same level of resolution become available.

In conclusion, Nanocompore offers a versatile, robust and practical method to readily identify RNA modifications from Nanopore DRS experiments. We envisage that its adoption by the scientific community will help to shed light on the distribution and function of RNA modifications at high resolution, helping to reveal the currently hidden life of RNAs.

## 4 Methods

### 4.1 Cell culture and KD/KO experiments

MOLM13 cells were cultured in RPMI1640 (Invitrogen) supplemented with 10% FBS and 1% penicillin/streptomycin/glutamine. Conditional knockdowns (KD) using METTL3-, DKC1-targeting or scrambled shRNAs were performed as previously described 8. For lentivirus production, 293T cells were transfected with PLKO.1 lentiviral vector containing the shRNA sequences (**Table S5**), together with the packaging plasmids psPAX2 (Addgene Plasmid #12260), and VSV.G (Addgene Plasmid #14888) for METTL3 KD or Pax2 (Addgene Plasmid #35002) and pMD2.G (Addgene Plasmid #12259) for DKC-1 KD experiments, at a 1:1.5:0.5 ratio, using Lipofectamine 2000 reagent (Invitrogen) according to the manufacturer’s instructions. Supernatant was harvested 48 and 72 h after transfection. 1×10^6^ cells and viral supernatant were mixed in 2ml culture medium supplemented with 8μg/ml polybrene (Millipore), followed by spinfection (60 min, 900g, 32°C) and further incubated overnight at 37°C. The medium was refreshed on the following day and the transduced cells were cultured further. MOLM13 cells (5×10^5^) were infected using PLKO-TETon-Puro lentiviral vectors expressing shRNAs. 24h after infection, the cells were replated in fresh medium containing 1 μg/ml of puromycin and kept in selection medium for 7 days. shRNA expression was induced by treatment with 200 ng/ml doxycycline for 4 days for METTL3 KD and 8 days for DKC1 KD experiments. For METTL3 knock-out (KO) experiments, lentiviruses were produced in HEK293 cells using ViraPower Lentiviral Expression System (Invitrogen) according to manufacturer’s instructions. MOLM13 cells stably expressing Cas9 were transduced with lentiviral gRNA vectors expressing either empty or METTL3 gRNAs (**Table S5**) and selected with puromycin from day 2 to day 5. At day 5 posttransduction, the cells were suspended in fresh medium without puromycin. At day 6, cells were harvested for RNA extraction.

### 4.2 RNA purification and *in vitro* transcription

Total RNA was isolated from MOLM13 cells using the RNeasy midi kit (Quiagen) and polyA+ RNA was purified from 30μg total RNA using the Dynabeads mRNA Purification Kit (Thermo Fisher Scientific) according to the manufacturer’s instructions. For production of unmodified 7SK RNA, synthetic double stranded DNA template for *in vitro* transcription (IVT) was produced by hybridization of synthetic Megamer^®^ Single-Stranded DNA Fragments (IDT) containing the 7SK sequence downstream of a T7 promoter (**Table S6**). 500ng of double stranded DNA template were used in 20μl IVT reactions for 1h using the TranscriptAid T7 High Yield Transcription Kit (Thermo Fisher Scientific), following the manufacturer’s instructions. The RNA product was purified using the RNA Clean & Concentrator kit (Zymo Research).

### 4.3 miCLIP

miCLIP was performed in duplicates with RNA isolated from wild type and METTL13 KO MOLM13 cells. The protocol is conceptually related to the original m6A miCLIP protocol^33^, but uses total RNA as input and follows a more recent variant of iCLIP protocol^34^. 4μg of total RNA were fragmented with RNA fragmentation reagents (ThermoFisher) following the manufacturer’s instructions. Fragmented RNA was then incubated with 2.5μg anti-m6A antibody (Abcam, ab151230) in IP buffer (50mM Tris-HCl pH 7.4, 100mM NaCl, 0.05% NP-40) at 4°C for 2 hours, in rotation. Subsequently, the solution was placed in 6-well plates on ice and irradiated twice with 0.3 J cm-2 UV light (254 nm) in a Stratalinker crosslinker. 30μl protein G beads (Dynabeads) per sample were washed twice with IP buffer and then incubated with the RNA-antibody solution at 4°C for 1.5 hours, in rotation. After the IP, the RNA-antibody-beads complexes were washed twice with High-Salt Wash buffer (50mM Tris-HCl pH 7.4, 1M NaCl, 1mM EDTA, 1% Igepal CA-630, 0.1% SDS, 0.5% sodium deoxycholate), once with IP buffer and once with PNK Wash buffer (20mM Tris-HCl pH 7.4, 10mM MgCl2, 0.2% Tween-20). The beads then proceeded to 3’ dephosphorylation and the rest of the iCLIP protocol. The 3’ adapters for on-bead ligation carry the sequences found in **Table S1**. Samples were mixed after the adapter removal step. Following the SDS-PAGE gel, the membrane was cut from 45kDa to 185kDa and RNA was extracted. The following sequence of the RT primer was used: /5Phos/ WWW CGTAT NNNN AGATCG-GAAGAGCGTCGTGAT /iSp18/ GGATCC /iSp18/ TACTGAACCGC. cDNA libraries were sequenced with single end 100bp reads on Illumina HiSeq4000.

### 4.4 Nanopore direct-RNA sequencing (DRS)

RNA sequencing was performed following the instruction provided by Oxford Nanopore Technologies (Oxford, UK), using R9.4 chemistry flowcells (FLO-MIN106) and direct-RNA chemistry sequencing kits (SQK-RNA001 or SQK-RNA002). For polyA+ transcriptome sequencing, we followed the conventional DRS protocol using the provided polyT (RTA) adapter. For the targeted sequencing, we ordered custom reverse transcription adapters complementary to the 3’end of 4 selected noncoding RNAs, and followed the sequencespecific DRS protocol (**Table S2**). For library preparation, we used 500ng of unmodified synthetic 7SK RNA using the adapter complementary to the 3’end of 7SK. To reduce sequencing costs for the DKC1 KD datasets, we multiplexed multiple conditions using the 4 barcodes supported by Poreplex (**Table S3**) and used 3μg of total RNA per sample for library preparation. The layout of the sequences used for targeted sequencing are represented in **Sup. Fig. S6**, page 29.

## 5 Data and computational methods

All the computational methods and scripts used for this paper are available in the following Github repository: https://github.com/tleonardi/nanocompore_paper_analyses. The direct RNA datasets generated for this study can be obtained from ENA (https://www.ebi.ac.uk/ena/browser/home) with the following accession number id: PRJEB35148. For this study we used the following Human reference files all obtained from Ensembl:

- Genome reference: Human genome assembly GRCh38.p12
- Annotation reference: Ensembl Gene build release-97

### 5.1 *In silico* simulated datasets

#### 5.1.1 Unmodified RNA model

We used an *in vitro* transcribed human RNA DRS dataset released by the Nanopore WGS consortium as a ground truth for non-modified RNA bases (https://github.com/nanopore-wgs-consortium/NA12878).

This dataset contains all possible 5-mers on average 58,307 times. The reads were aligned on gencode release 28 human reference transcriptome with Minimap2v 2.14 and we realigned the signal to the reference sequence using Nanopolish eventalign v0.10.1 followed by NanopolishComp Eventalign_collapse v0.5. Next, we collected the median intensity and dwell time data for each 5mers and tried to fit 44 different distributions. We selected distributions minimising the sum of square root error for all kmers between the observed and modeled data. In addition, we also based our selection on the possibility to easily change the parameters of the distributions to simulate the presence of modifications. We selected the Wald distribution and the Logistic distribution for dwell time and median intensity, respectively. Finally, we generated a model file containing the parameters of the observed and model distributions for each 5-mer. The up-to-date model file is distributed with Nanocompore. The detailed analysis is available in the following Jupyter notebook: https://github.com/tleonardi/nanocompore_paper_analyses/blob/master/in_silico_dataset/01_IVT_Kmer_Model.ipynb.

#### 5.1.2 Simulated reference sequence

We generated a set of *in silico* reference sequences. In order to maximise the sequence diversity and kmer coverage we used a “guided” random sequence generator. In brief, the sequences are generated base per base using a random function, but the program keeps track of the number of times each kmer was already used. The sequence is extended, based on a random function with a weighted probability for each kmer inversely proportional to their occurrence in the sequences already generated. This ensures that all kmers are represented as uniformly as possible, but it leaves some space to randomness. We generated a set of 2000 sequences 500 bases long each maximising the 9-mers coverage. We excluded any homopolymers longer than 5 bases, as they are likely to be miscalled in nanopore data. Kmer coverage in the final sequence set are summarised in **Table S4**. The detailed analysis is available in the following Jupyter notebook: https://github.com/tleonardi/nanocompore_paper_analyses/blob/master/in_silico_dataset/02_Random_guided_ref_gen.ipynb.

#### 5.1.3 Simulated modified and unmodified datasets

Nanocompore comes with a companion tool called SimReads which can generate simulated read data based on a fasta reference and a kmer model file. Essentially, SimReads walks along the reference sequence and generates intensity and dwell time values corresponding to each 5-mers. To do so, it uses a probability density random generator using the kmer model values (location and scale) bounded by the extreme observed values. This tool can also offset the model mean by a fraction of the distribution standard deviation to simulate the effect of RNA modifications. This can be done for all the reads or only on a subpopulation of reads. SimReads generates files similar to the output of NanopolishComp EventalignCollapse. This means that the datasets can be directly used as input for NanoCompore SampComp. Using Nanocompore v1.0.0rc3 with the previously described simulated reference sequence set we generated 144 *in silico* datasets with various amplitude of modification of the median signal intensity and the dwell time (0, 1, 2, 3 and 4 standard deviation) as well as different fractions of modified reads (10%, 25%, 50%, 75%, 90% and 100%). All the datasets were simulated in duplicate with a uniform coverage depth of 100 reads. The detailed analysis is available in the following Jupyter notebook: https://github.com/tleonardi/nanocompore_paper_analyses/blob/master/in_silico_dataset/03_Simulated_dataset_gen.ipynb.

#### 5.1.4 Analysis of simulated datasets

We compare the 144 datasets containing simulated modifications against the reference dataset generated from the unmodified model with Nanocompore v1.0.0rc3 (See Nanocompore section after). The analysis was performed with all the statistical methods supported by Nanocompore using a sequence context of 2 nucleotides (https://github.com/tleonardi/nanocompore_paper_analyses/blob/master/in_silico_dataset/04_nanocompore.sh). The result database was subsequently parsed and the predicted modified sites were compared with the position of the known simulated positions. A hit was considered true positive (TP) when we found a significant p-value within 3 nucleotides of a known modified position. A significant hit outside of this window was counted as a false positive (FP). Finally, we plotted the Receiver Operating Characteristic (ROC) curves corresponding to the TP rate compared with the FP rate for every Nanocompore comparison performed (https://github.com/tleonardi/nanocompore_paper_analyses/blob/master/in_silico_dataset/05_calc_roc.sh).

### 5.2 Direct-RNA datasets analysis

#### 5.2.1 Demultiplexing

For targeted sequencing of ncRNAs in the DKC1 KD vs WT experiment, we multiplexed several conditions together using custom barcodes as described in the section “Nanopore direct-RNA sequencing” and outlined in **Table S3**. The datasets were demultiplexed using Poreplex v0.4 with default parameters (https://github.com/hyeshik/poreplex/tree/v0.4). We were able to assign 23% of the reads to one of the 4 barcodes used.

#### 5.2.2 Data preprocessing

All the datasets were preprocessed using an automated analysis NextFlow pipeline, before running Nanocompore (https://github.com/tleonardi/nanocompore_pipeline). Raw reads FAST5 files were basecalled with ONT Guppy v3.1.5 and the basecalled reads were saved in FASTQ format. A post-basecalling quality control was performed with pycoQC (v2.2.4)^24^ to verify the consistency of the sequencing runs. A transcriptome reference FASTA file was created from the annotation BED file and genome FASTA file with Bedparse (v0.2.2)^35^. Reads were then aligned on the transcriptome reference with Minimap2 (v2.16)^36^ in unspliced mode (-x map-ont). The resulting aligned reads were filtered with samtools (v1.9)^37^ to keep only primary alignments mapped on the forward strand (-F 2324) and the raw signal was realigned on reads using Nanopolish eventalign (v0.11.1)^19^. Finally, the data was processed by NanopolishComp Eventalign_collapse (v0.6.2)^38^ to generate a random access indexed tabulated file containing realigned median intensity and dwell time values for each kmer of each read.

#### 5.2.3 Synthetic m6A data preprocessing

We obtained a dataset corresponding to an *in vitro* oligomer containing the m6A-modified and unmodified GGACU motifs ligated to a large carrier RNA^16^. Briefly, the fast5 files were basecalled with Guppy (v3.1.5) and the resulting sequences mapped to the reference with minimap2 (v2.16). The raw signal was then realigned to the mapped sequences with Nanopolish eventalign (v0.11.1) and the resulting eventalign file was split into two separate files respectively containing the modified and unmodified UGAGGACUGUA subsequence. After splitting, the resulting files were processed with NanopolishComp Eventalign_collapse and Nanocompore as described for the other datasets. The code for this analysis is available at the following URL: https://github.com/tleonardi/nanocompore_paper_analyses/tree/master/synthetic_oligo_timp/scripts.

### 5.3 Signal comparison with Nanocompore

Nanocompore is a Python3 package dedicated to comparative analysis of DRS nanopore sequencing raw signal in order to identify potential RNA modifications sites. Signal analysis and complex statistical tests are generally resource-intensive, but Nanocompore takes advantage of a multiprocessing architecture to process transcripts in parallel and has a relatively small memory footprint. Nanocompore requires at least 1 indexed tabulated file generated with NanopolishComp Even-talign_collapse for each of the 2 conditions to compare. The program will run with a single replicate per condition, but we recommend at least 2 to take full advantage of the advanced statistical framework. The analysis flow is divided in three steps: 1) white-listing of transcripts with sufficient coverage, 2) parallel processing and statistical testing of transcripts position per position, 3) post-processing and saving.

#### 5.3.1 Transcripts whitelisting

In order to reduce the computational burden, Nanocompore first filters out transcripts with insufficient coverage. This is achieved by a rapid tally of reads mapped per transcripts followed by selection of transcripts having at least 30 reads mapped in all of the samples provided. Users can modify the threshold but the default value allows to get reproducible results. Optionally, one can provide a custom list of transcripts to include or exclude.

#### 5.3.2 Statistical analysis

White-listed transcripts are processed in parallel to take advantage of multi-threaded architecture. First, the data corresponding to the reads mapped on each transcript is loaded in memory and transposed in the transcript space in a position-wise fashion. The current implementation of Nanocompore only uses the median signal intensity and the scaled log_10_ transformed dwell time, but the framework is flexible enough to aggregate more variables, such as the error rate or additional Nanopolish HMM states. The 2 experimental conditions are compared positions per position using a range of statistical tests. We included 3 robust univariate pairwise statistical methods: Kolmogorov-Sirmrnov (KS), Mann-Whitney U (MW) and t-test. Those tests are performed independently on the median intensity and the dwell time. We also implemented a Gaussian mixture model (GMM) clustering-based method. For a given position we fit a bivariate 2 components GMM to all the data points observed (x=median intensity, y=dwell time), irrespective of the sample label. We then assign each data point to one of the two clusters and test for differences in the distribution of reads between clusters across conditions. To this purpose, testing is implemented in two ways: 1) we fit a Logit model to the data using the formula predicted_cluster~1+sample_label and report the coefficient’s p-value. 2) We do a one-way ANOVA test comparing the log odds of data points belonging to cluster one between the two conditions. After testing, it’s optionally also possible to aggregate the p-values of neighbouring kmers to account for the fact that modified bases affect the signal of multiple kmers. To this end, and due to the fact that neighbouring p-values are non-independent, we implemented in python a method to approximate the distribution of the weighted combination of non-independent probabilities^23^. The combined p-values are computed all along the sequence using a sliding window of a given length. This method greatly reduces the prediction noise (false positive rate) at the expense of spatial resolution, while giving more weight to sites for which the effect of RNA modifications on the signal is spread over several kmers.

#### 5.3.3 Post-processing, saving and data exploration with Nanocompore interactive plotting API

Results generated by the statistical module are collected and written in a simple key/value GDBM database. Although this data structure has limitation in terms of portability and concurrent access, it is natively supported by python and allow to store complicated data structures. For each test previously performed p-values are temporary loaded in memory and corrected for multiple tests with the Benjamini-Hochberg procedure. Users can then obtain a tabulated text dump of the database containing all the statistical results for all the positions in the transcripts space or a BED file with the positions of significant hits found by Nanocompore converted in the genome space. Finally, we provide a convenient python wrapper over the GDBM database, allowing users to interactively access simple high level functions to plot and export the results (https://nanocompore.rna.rocks/demo/SampCompDB_usage/). The wrapper was initially developed for Jupyter but can essentially work with any python IDE. At the time of publication the wrapper allows to generate 6 different types of publication ready plots for a given transcript including (1) the distribution of p-values, (2) the distribution of signal intensity and dwell time, (3) the overall coverage per sample, (4) the nanopolish HMM states, (5) the kernel density of the signal and dwell time for a specific position and (6) the sharkfin plot of the p-values compared with Log Odds Ratio (for the GMM method). The API, will be progressively extended in the future.

### 5.4 Downstream analyses

The code for all generic analyses, plots and metrics is available at https://github.com/tleonardi/nanocompore_paper_analyses/. The transcript intersection plot for the MOLM13 polyA dataset had been generated with UpsetR^39,40^.

#### 5.4.1 Metagene m6A coverage

The metagene m6A coverage analysis was done considering all nanocompore sites with GMM logit p-value<0.01. The plot was produced with GuitarPlot using m6A sites in genome-space (from the BED files produced by Nanocompore) and the Bioconductor TxDb.Hsapiens.UCSC.hg38.knownGene packages as txdb option.

#### 5.4.2 Motif enrichment analysis of m6A sites

For the motif enrichment analysis of m6A sites identified by Nanocompore analysis of METTL3 KD, we extracted the sequence of all kmers tested by Nanocompore and having a p-value<0.5 (GMM-logit). The sequences were then sorted by p-value and analysed with Sylamer for the identification of over-represented words, using a word size of 5 and a growth parameter of 100. The Sylamer results were then imported in R for plotting. For visualisation purposes, the final plot only reports the lines for the top 100 motifs with the greatest area under the sylamer curve, with the top one represented in colour.

#### 5.4.3 Single molecule identification of m6A sites

To assign an m6A probability at A652, A1324 and A1535 for each read covering the β-actin transcript, we developed a dedicated post-processing script available at https://github.com/tleonardi/nanocompore_paper_analyses/m6acode/parse_sampcomdb.py. Briefly, for each of the three positions of interest, we extract the GMM model saved in sampCompDB, and for each read we then predict the probability that it belongs to each of the two clusters. To define which of the two clusters corresponds to m6A modified reads, we consider which of the two clusters has negative log odds of data points belonging to it in the KD condition (i.e., we consider which of the two clusters shrinks in the KD condition). To test the independence of the methylation events at these three sites, we performed a chi-squared test of independence comparing the expected number of molecules for each of the 8 combinations of modifications to the observed number of molecules. The results reported are obtained using a probability threshold of 0.75 (as predicted by the GMM) to consider a read as methylated. However, to ensure robustness of these results, the chi-squared test was repeated for all thresholds between 0.1 and 1 (0.05 steps) and p-values were adjusted accordingly using the Benjamini–Hochberg procedure. Adjusted p-values were >0.39 for all thresholds used.

#### 5.4.4 7SK structures

The 7SK multiple alignments and consensus secondary structure were obtained from Rfam (RF00100). Secondary structure plots were produced with R2R^34,41^ and a custom python script to annotate p-values as color shading (available at https://github.com/tleonardi/nanocompore_paper_analyses/blob/master/ncRNAs_structures/create_annotations.py).

#### 5.4.5 miCLIP analysis

miCLIP data and corresponding input data was analysed using the iMaps web server (https://imaps.genialis.com/). Briefly, raw reads were demultiplexed and trimmed (for adaptors and quality), before being mapped to a tRNA and rRNA index using STAR (v2.4.0.1)^42^. Unmapped reads were then mapped to GRCh38 GENCODE primary assembly, using GENCODE annotation v30. STAR parameter – alignEndsType Extend5pOfRead1 was used to ensure no soft clipping of cDNA start sites. PCR duplicates were removed based on unique molecular identifier (UMI) and mapping position. cDNA start −1 positions were taken as crosslink sites. To calculate miCLIP coverage of Nanocompore sites, the resulting BAM files were converted to transcriptome space. Briefly, the BAM files produced from the iMaps pipeline where filtered to only retain uniquely mapping reads and then converted to fastq format with Samtools. The reads were then mapped with bowtie2 to an index generated from the sequences of all the 752 transcripts analyzed by Nanocompore. The resulting BAM files were then loaded into R, the start positions converted to GRanges with BAM2GRanges and normalised using the TMM method as implemented in csaw^43^. The binned coverage was calculated using the featureScores function of the Repitools package^43,44^ with a bin size of 50nt. To test the null hypothesis of no difference between WT and KO, we calculated the mean coverage of the bins within 100nt of a Nanocompore site and applied the Mann-Whitney U test. For these analyses only Nanocompore sites with GMM-logit p-value <0.01 were considered and sites less than 5nt apart were merged.

## 6 Acknowledgements and conflict of interest

We would like to thank the DNA Pipelines Research and Development group at the Wellcome Sanger Institute for their support and help, in particular Kim Judge. We also thank Andrew Bannister, Mattia Pelizzola, Mattia Furlan, Michael Harbour and Jack Monahan for their constructive comments and for proofreading the manuscript.

EB is a paid consultant of ONT. TK is a co-founder of STORM Therapeutics Limited. PPA is a Research Scientist at STORM Therapeutics Limited.

## Supplementary Figures and Tables

**Figure 1:**
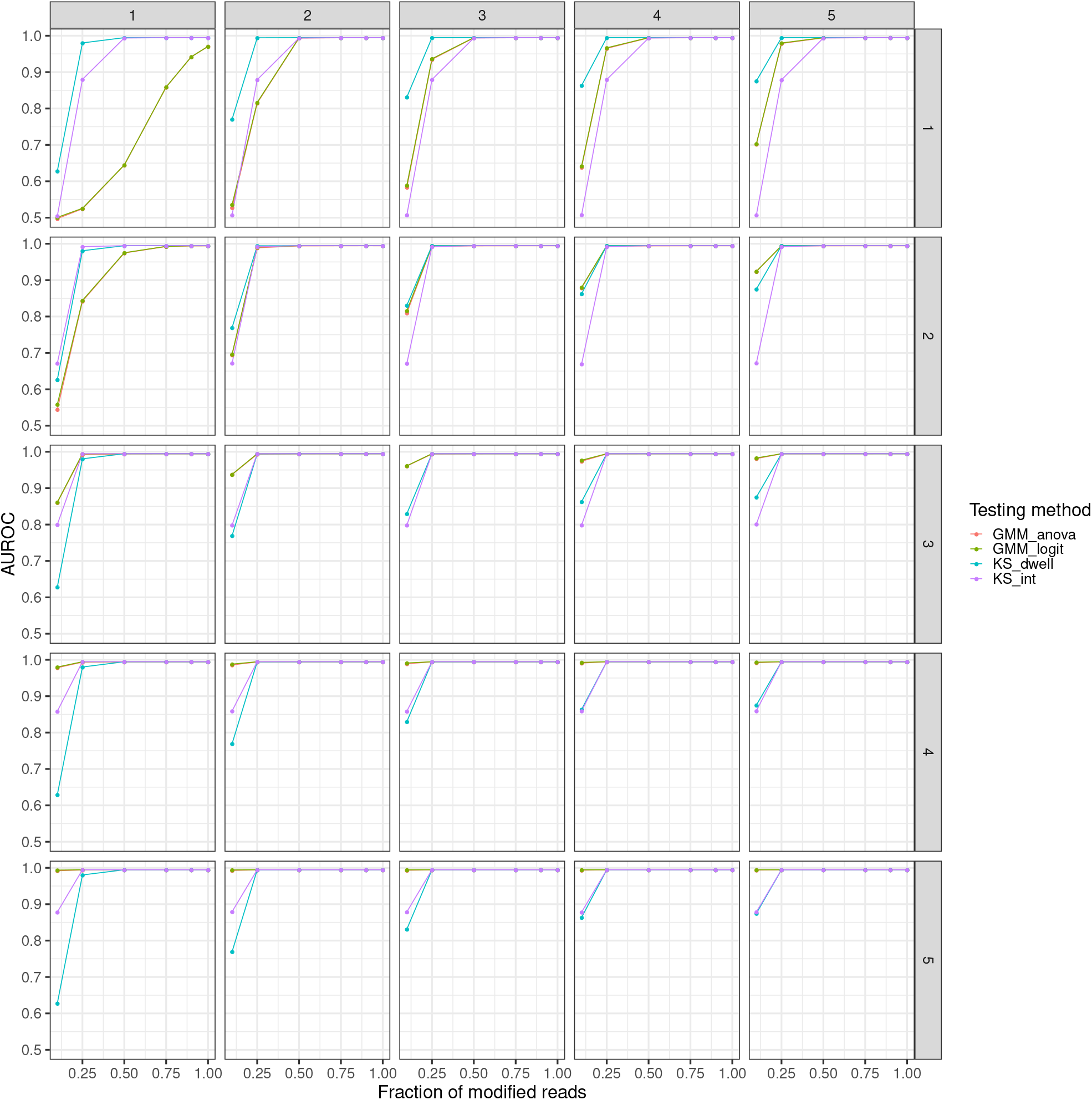
Benchmarks on *in silico* data. Plots showing the Area Under the ROC curve (AUROC, y-axis) obtained with Nanocompore on *in silico* generated data at varying fractions of modified reads (x-axis).

**Figure 2:**
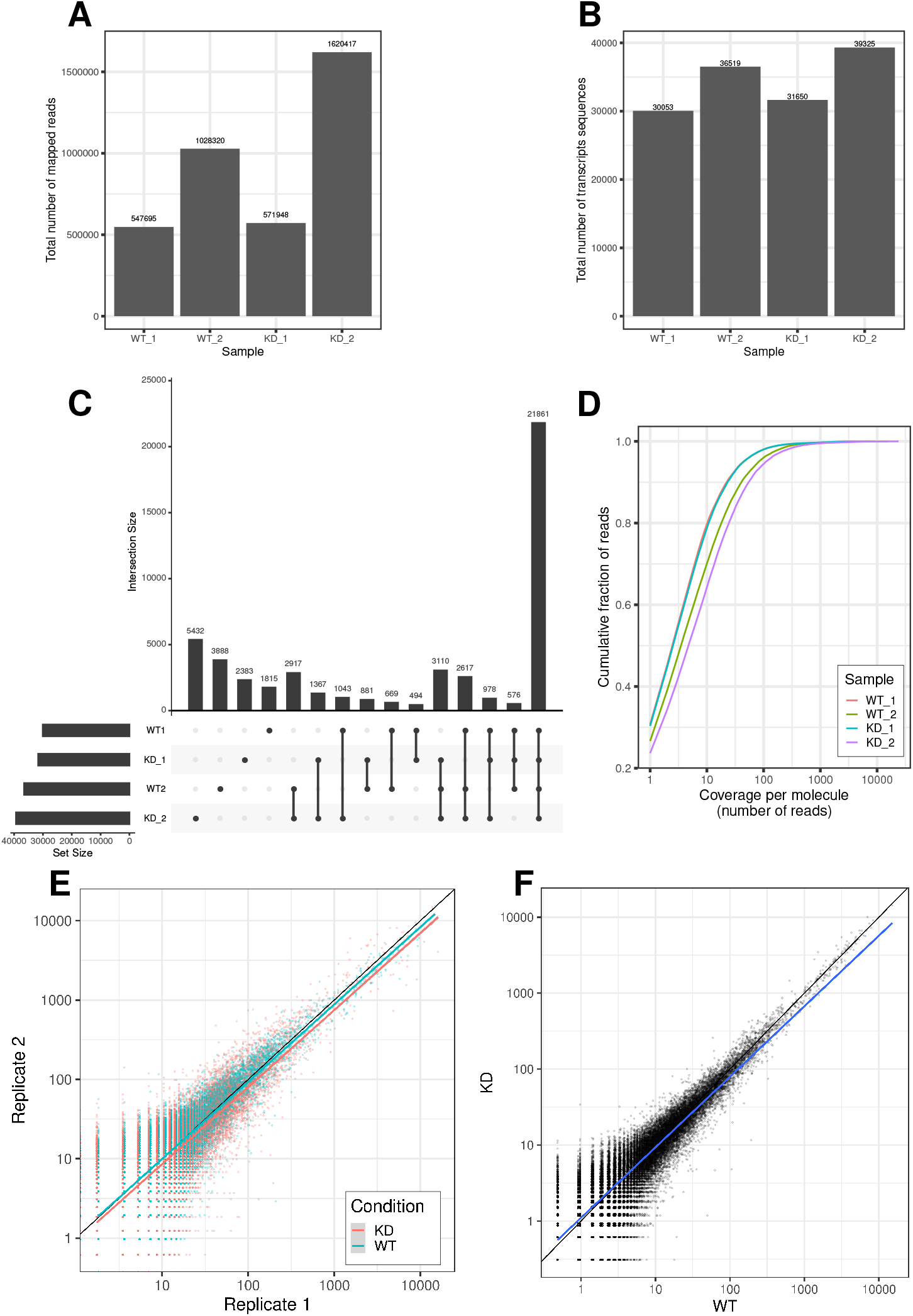
m6A profiling in the polyA+ transcriptome. **A:** Bar chart showing the total number of mapped reads in each sample. Overall total number of reads: 3,768,380 (average 942,095 per sample). **B:** Bar chart and intersections plots showing the total number of sequences transcripts in each sample. Average: 34,386.75 distinct transcripts per sample **C:** Bar chart and intersections plots showing the total number of sequences transcripts in each sample and in every combination of samples. **D:** Cumulative fraction of reads mapping to transcripts with increasing degree of total coverage. For example, the point (10,0.6) on the purple line indicates that in the KD2 sample ~60% of the reads map to transcripts with coverage of 10x or lower. **E:** Scatter plot showing the correlation in transcript abundance between replicates. R^2^ of 0.757 and 0.937 for WT and KD respectively. **F:** Scatter plot showing the correlation in transcript abundance between WT and KD after averaging replicates. R^2^=0.969.

**Supplementary Table 1:**
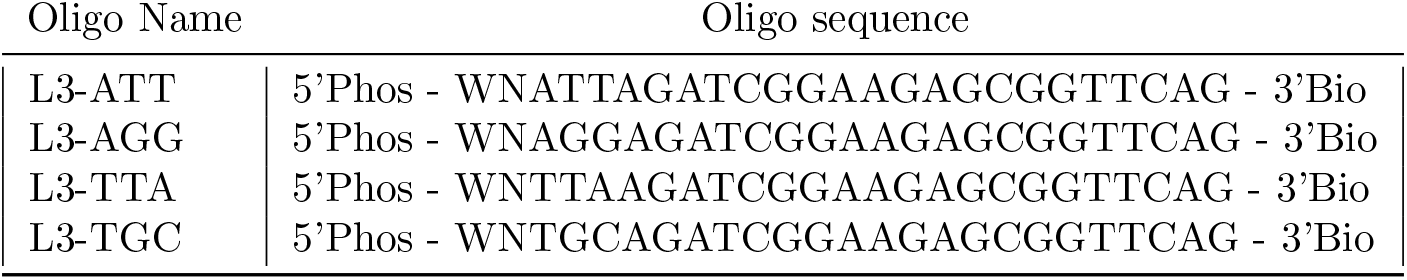
Sequence of miCLIP barcoded adapters

**Supplementary Table 2:**
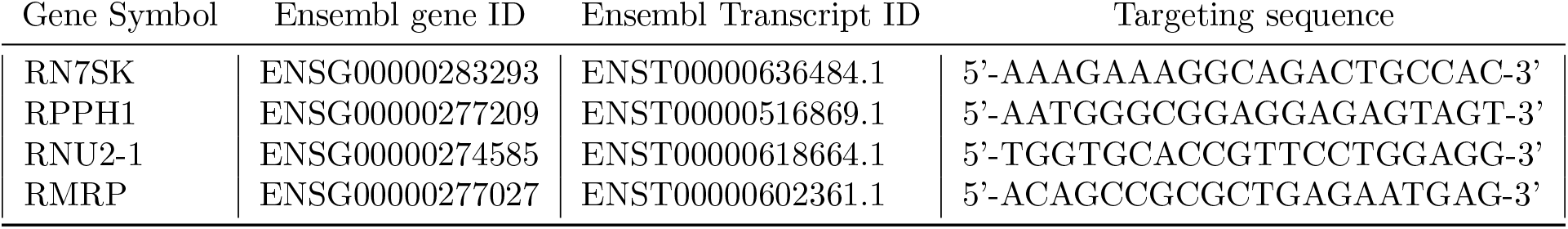
Targeted ncRNA

**Supplementary Table 3:**
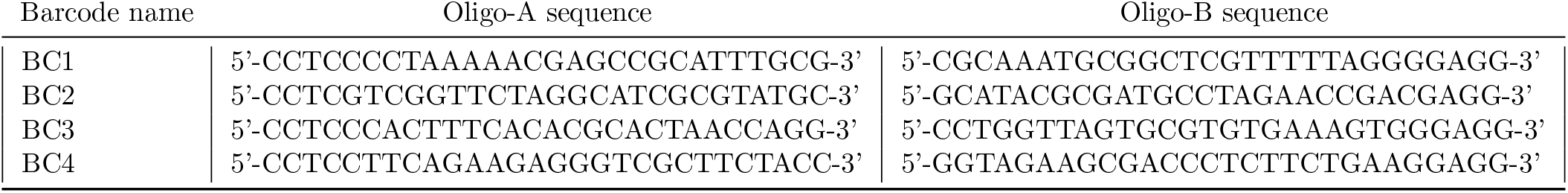
Poreplex compatible barcoding sequences

**Supplementary Table 4:** Coverage of kmers (5 to 11 mers) found in the *in silico* generated reference sequences. The sequence set was designed to optimise the 9-mers content. The expected kmer count excludes the sequences containing homopolymers longer than 5 bases which were intentionally avoided when designing the reference sequence.

| Order | Median occurrences | Mean occurrences | Min occurrences | Max occurrences | kmers found | kmers expected | Percent kmers found |
| --- | --- | --- | --- | --- | --- | --- | --- |
| 5-mers | 970 | 968 | 695 | 1023 | 1024 | 1024 | 100 |
| 6-mers | 242 | 241 | 214 | 275 | 4092 | 4092 | 100 |
| 7-mers | 60 | 60 | 45 | 89 | 16356 | 16356 | 100 |
| 8-mers | 15 | 15 | 8 | 26 | 65376 | 65376 | 100 |
| 9-mers | 4 | 3 | 0 | 9 | 261268 | 261312 | 99.98 |
| 10-mers | 1 | 1 | 0 | 6 | 706080 | 1044480 | 67.6 |
| 11-mers | 1 | 1 | 0 | 4 | 913965 | 4174848 | 21.89 |

**Figure 3:**
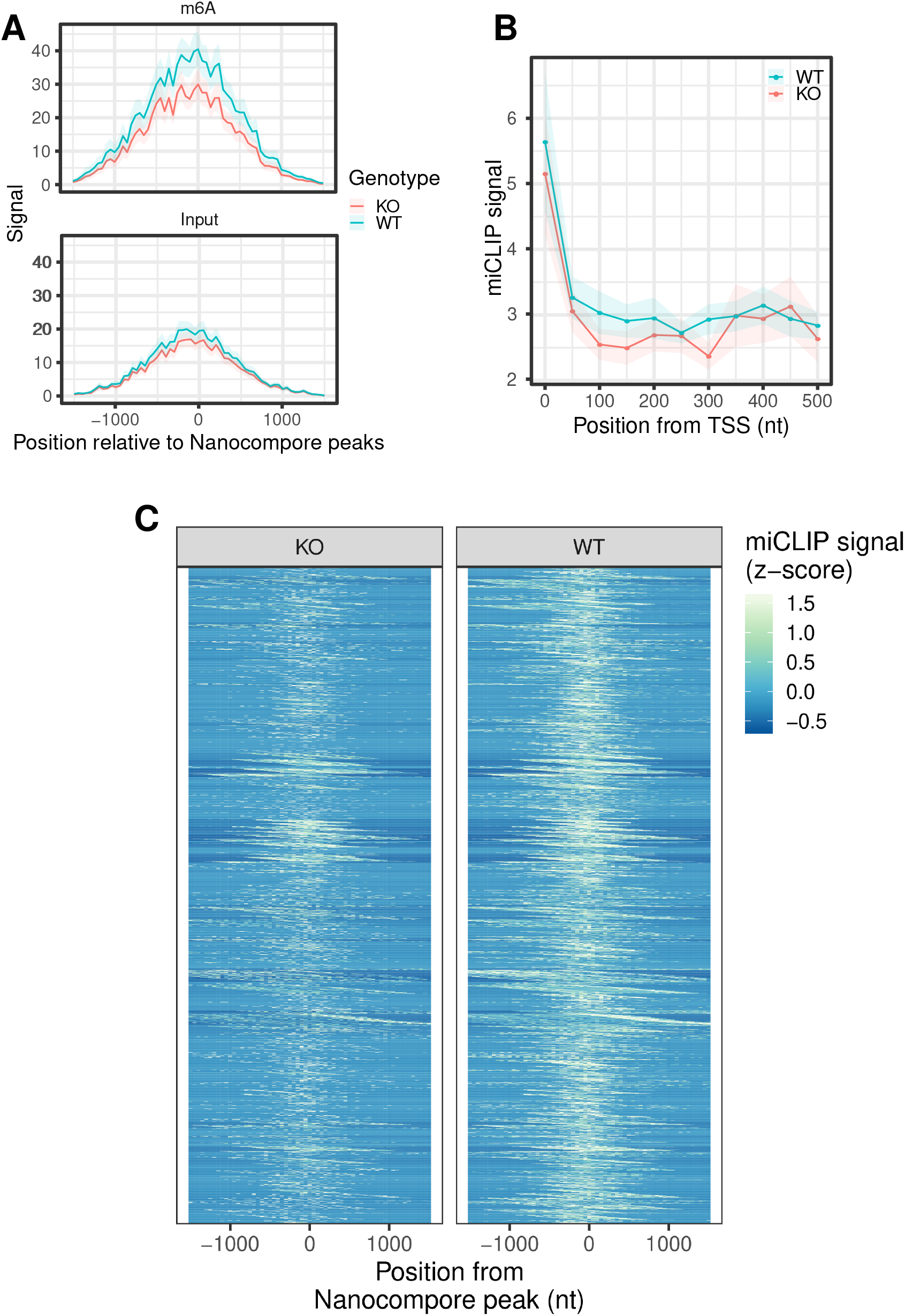
miCLIP validation of Nanocompore m6A sites. **A:** m6A miCLIP coverage of significant Nanocompore sites (GMM logit p-value<0.01). Coverage calculated in transcriptome-space. The y-axis shows the mean signal across sites of the average coverage (counts per million) in 4 replicates for the WT condition and 2 replicates for the KO condition. The shaded area shows the standard error of the mean across sites. **B:** Same as in A but showing the coverage of the transcriptional start sites (TSSs) of transcripts with at least one significant Nanocompore site. **C:** Heatmap showing the z-score normalised mean coverage around significant Nanocompore sites. For visualisation purposes the color scale was saturated at the 95th percentile of distribution of z-scores.

**Supplementary Table 5:**
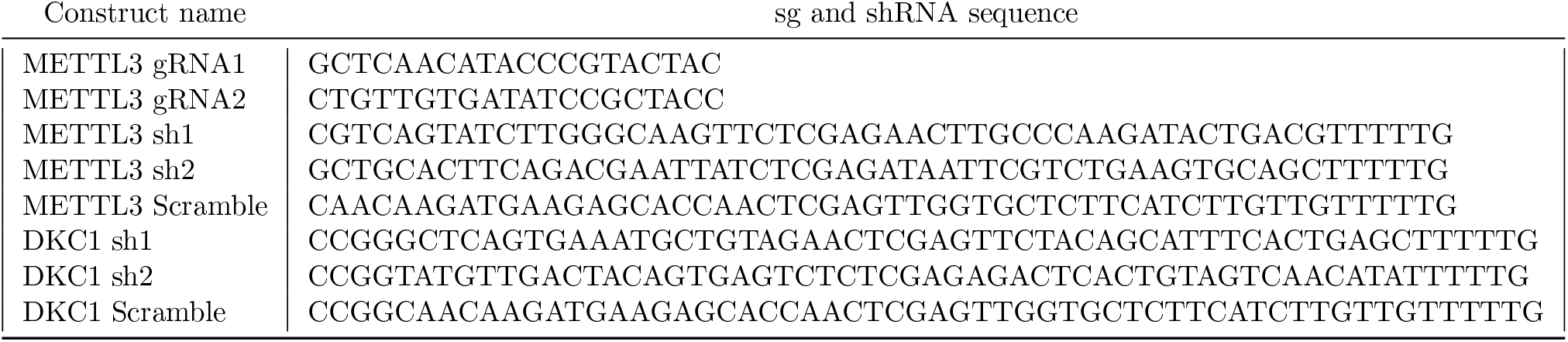
Sequences of sgRNAs and shRNAs used for KD and KO experiments.

**Supplementary Table 6:**
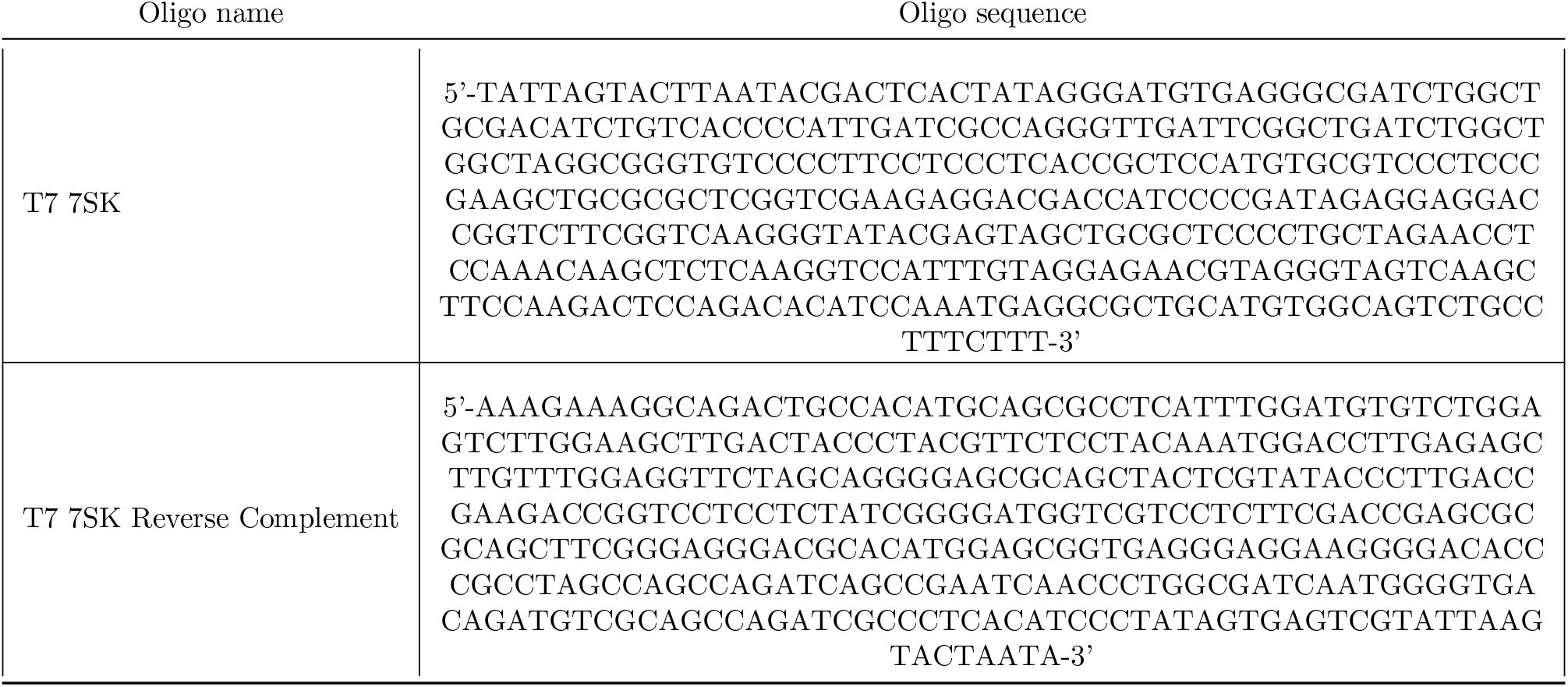
DNA oligo sequences used to produce double stranded DNA template for 7SK *in vitro* transcription. The T7 promoter is in the first and last 28 nucleotides of the forward and reverse complement sequence respectively.

**Figure 4:**
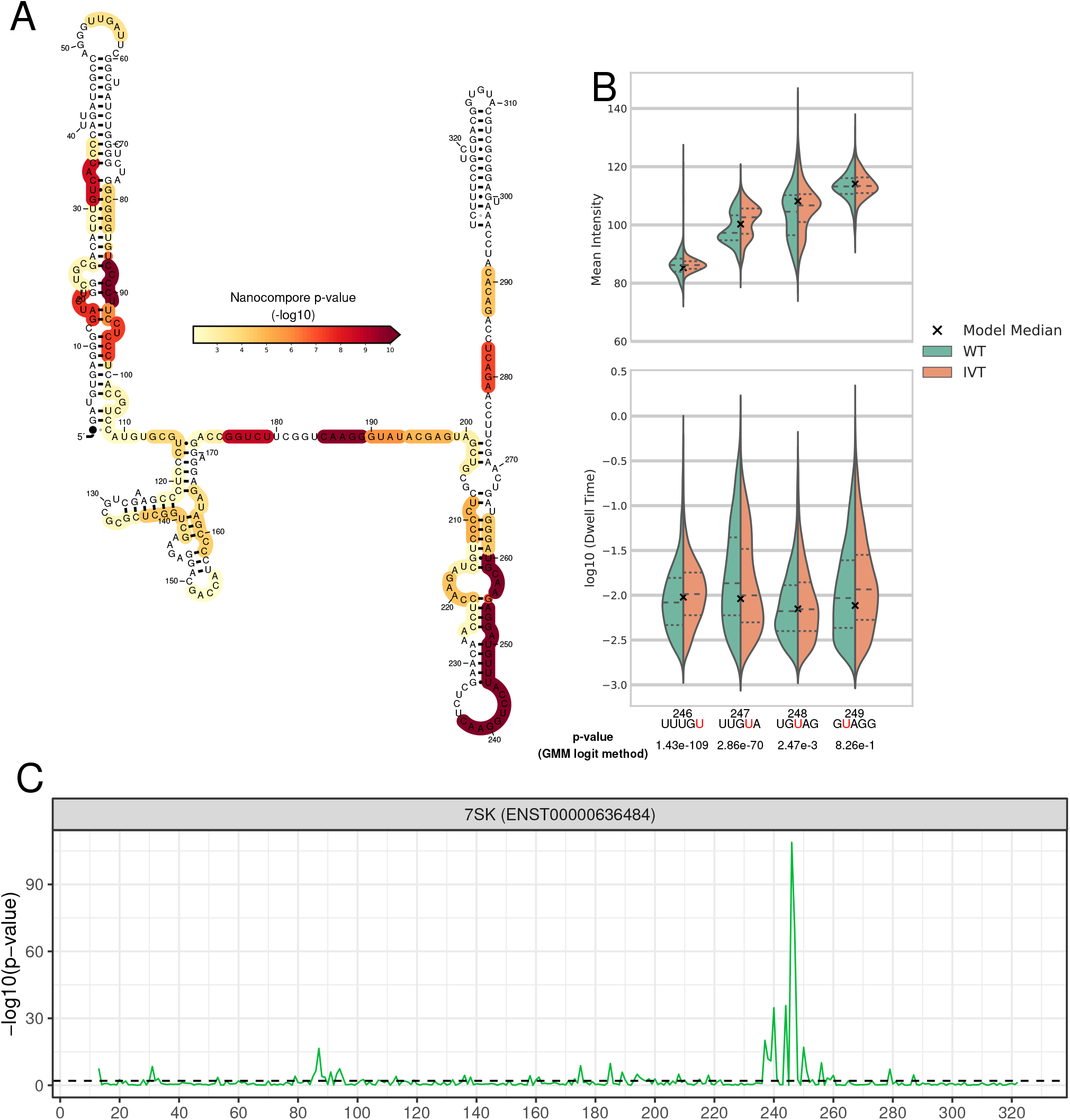
Modification profile of 7SK from the analysis on an IVT sample. **A:** Secondary structure of 7SK with the Nanocompore IVT vs WT p-value (GMM-logit) overlaid as a color scale. For each nucleotide the color indicates the lowest p-value among those of the 5 kmers that overlap it. Only p-values<0.01 are shown in color. **B:** Violin plots showing the distributions of median intensity (top) and scaled log10 dwell time (bottom) for kmers encompassing the known pseudouridine site U250. the Hexim1 binding sites and neighbouring kmers. **C:** RNA modification profile of 7SK, showing the Nanocompore GMM-logit p-value (y axis, −log10) across the transcript length. All coordinates refer to the first nucleotide of each kmer relative to ENST00000636484.

**Figure 5:**
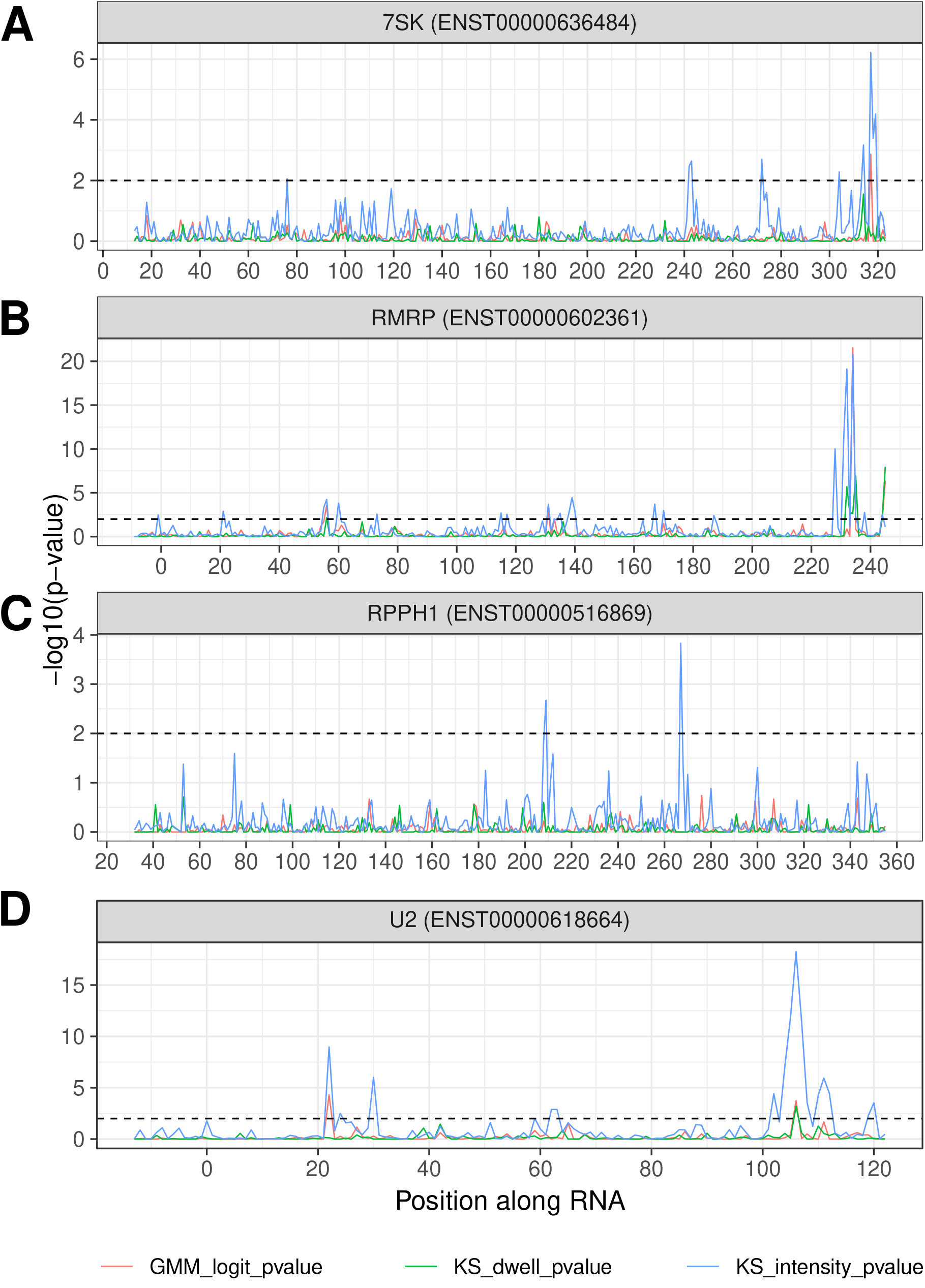
Pseudouridine profile of ncRNAs. **A-D:** DKC1-dependent pseudouridine profiles 7SK, showing the Nanocompore p-values (y-axis, DKC1 KD vs control, −log10) across the transcript length for 7SK (A), RMRP (B), RPPH1 (C) and U2 (D). All coordinates refer to the first nucleotide of each kmer.

**Figure 6:**
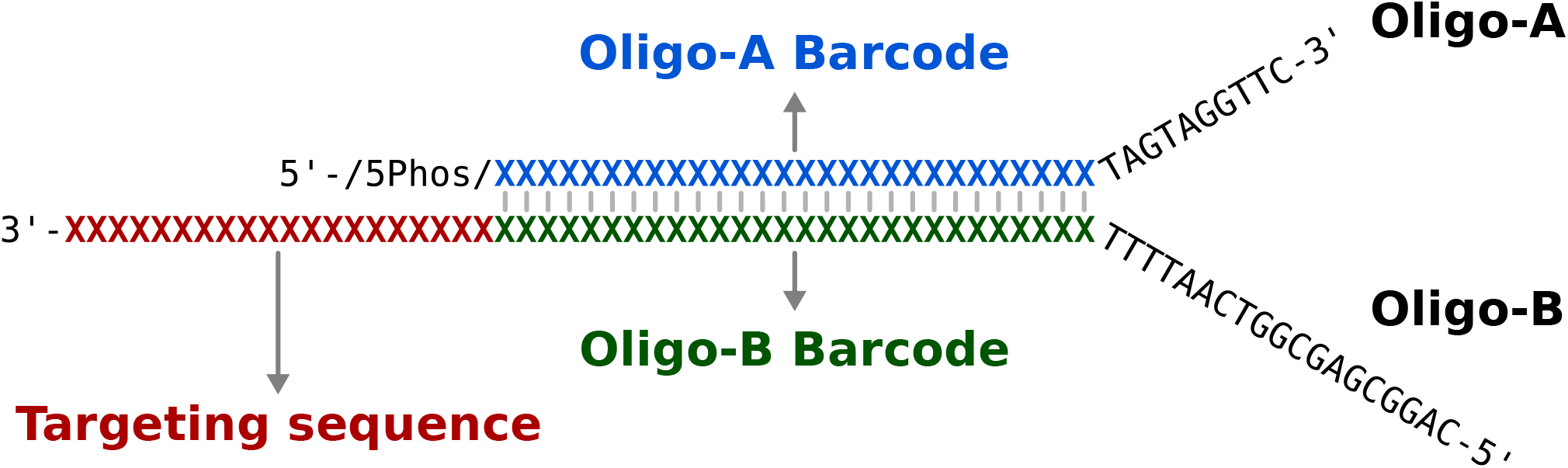
Sequence of the custom RT/sequencing adapter used for targeted sequencing.

